# Altered frequency of CD24^high^CD38^high^ transitional B cells in patients with cardiac involvement of chronic Chagas disease

**DOI:** 10.1101/684589

**Authors:** Magalí C. Girard, Gonzalo R. Acevedo, Micaela S. Ossowski, Paula B. Alcaráz, Marisa Fernández, Yolanda Hernández, Raul Chadi, Karina A. Gómez

## Abstract

The cardiomyopathy developed by patients with chronic Chagas disease (CCD), one of the most severe consequences of *T. cruzi* infection, is mainly associated with an imbalance between an excessive inflammatory reaction and a defective immunomodulatory profile cause by host-parasite interaction. Despite the growing importance of the regulatory function of B-cells in many malignancies, few studies have addressed their immunosuppressive role in chronic Chagas disease. In this work, we tackled this issue by studying the proportion of different B cell subpopulations and their capacity to secrete IL-10 in individuals with distinct clinical forms of CCD. Seven-colour flow cytometry was performed to examine the peripheral blood B cell compartment in chronic Chagas disease (CCD) patients with and without cardiac manifestations (n=10 for each group) and non-infected donors (n=9). Peripheral blood mononuclear cells (PBMC) were incubated for 5h with PMA, ionomicyn and brefeldin A. According to the expression of markers CD19, CD24 and CD38, we showed an expansion of total B cell and transitional CD24^high^CD38^high^ B cell subsets in CCD patients with cardiac involvement compared to non-infected donors. Furthermore, although no differences were observed in the frequency of total IL-10 producing B cells (B10) among the groups, CCD patients with cardiac involvement showed a statistically significant increased proportion of naïve B10 cells and a tendency to an increased frequency of transitional B10 cells compared to non-infected donors. These findings suggest that immature transitional CD24^high^CD38^high^ B cells are greatly expanded in patients with the cardiac form of chronic Chagas disease and these cells retain their ability to secrete IL-10 compared to non-infected donors. Furthermore, the distribution of naïve, transitional and memory B cells inside the B10 cells followed the same pattern in chronic patients without cardiac involvement and non-infected individuals. Our work provides insight into the phenotypic distribution of regulatory B cell in CCD, an important step towards new strategies to prevent cardiomiopathy associated with *T. cruzi* infection.

## INTRODUCTION

Chagas disease, a serious health problem caused by the infection with the protozoan parasite *Trypanosoma cruzi*, comprises an acute and a chronic phase. During the latter, the disease may remain without any detectable symptoms for several decades, or progress toward cardiac or digestive forms, or even a combination of these alterations (1). The absence of any symptoms as well as the inflammatory mechanisms leading to tissue damage is attributed mainly to the immune response developed by the host against the parasite, and to the different means by which the parasite avoids it and persists. Several reports demonstrate a predominance of a humoral and cellular pro-inflammatory environment in cardiac patients, while an anti-inflammatory response seems to prevail in infected subjects without cardiac manifestation (2-4).

Regulatory B (Breg) cells are a specific B cell subset allocated to restrain the excessive inflammatory response accomplished in different immune-related pathologies, such as autoimmune and allergic diseases, malignancies, infections and solid organ transplantation (5, 6). Through IL-10 production, Breg cells have the capacity to suppress the function and proliferation of Th1, Th17 and follicular helper T (T_FH_) cells, increase polarization of T cells towards the regulatory T (Tregs) cell profile, repress the innate response by acting on antigen-presenting cells (dendritic cells, macrophages) and natural killer (NK and NKT) cells, decrease the production of IgG while inducing class switch towards IgG4, among others (7). In addition, Breg cells have also been shown to contribute to immune homeostasis by IL-10-independent mechanisms (8). However, and due to the absence of subset-specific membrane markers or transcription factors, IL-10 secretion is still the hallmark feature used to distinguish Breg from the rest of the B cells (9, 10). In humans, different B cell lineages that secrete this cytokine and have regulatory functions have been described, such as CD19^+^CD24^high^CD38^high^ (11), CD19^+^CD24^high^CD27^+^ (12), CD19^+^CD5^+^CD1d^+^ (13, 14), CD19^+^Tim-1^+^ (15, 16) and CD19^+^CD25^+^CD71^+^CD73^−^ cells (7), depending on the stimulation conditions and the markers used to identify them. Nowadays, the majority of the studies in human samples pinpoint the CD19^+^CD24^high^CD38^high^ immature transitional B cell population as the most representative phenotypic signature of Breg cells (17).

In the context of Chagas disease, B cells and their role as antibody secreting cells were amongst the first and most widely studied components of immunity against *T. cruzi*, but little is known about the irrelevance as antigen presenting cells, cytokine producers and immune modulators (3, 18). To date, only Fares *et al* (2013)(19), described that patients with chronic Chagas disease have an increased frequency of IL-10- and TGF-β-producing B cells in peripheral blood, both in basal state and upon *in vitro* stimulation with parasite lysate, hinting that these cells participate in the delicate balance between protection and pathogenesis.

Given that Breg cells have the ability to ameliorate exacerbated inflammatory responses, hampering the development of tissue damage while contributing to pathogen persistence, we sought to analyze the frequency and phenotypic distribution of B and B10 cells according to their expression of CD24, CD38 and CD27, in peripheral blood from patients with chronic Chagas disease with or without cardiac involvement.

## MATERIALS AND METHODS

### Subjects included and blood sample collection

EDTA-anti coagulated blood samples were collected from patients with chronic Chagas disease (CCD) and from non-infected donors after written consent was given, in accordance with the guidelines of the protocol approved by the Medical Ethics Committee of the Instituto Nacional de Parasitología “Dr. M. Fatala Chaben” and the Hospital General de Agudos “Dr. Ignacio Pirovano”. The study population included 30 subjects: 20 patients with CCD (10 with and 10 without cardiac manifestations) and 9 donors with non-reactive serological tests for Chagas disease (control group). *T. cruzi* chronic infection was determined, according to national and international guidelines, by at least two reactive tests (indirect immunofluorescence, enzyme-linked immunosorbent assay –ELISA-, indirect hemagglutination). All the three groups were age- and gender-matched (**Table 1**). The exclusion criteria included record of treatment with benznidazole or nifurtimox and presence of: systemic arterial hypertension, diabetes mellitus, thyroid dysfunction, renal insufficiency, chronic obstructive pulmonary disease, hydroelectrolytic disorders, alcoholism, history suggesting coronary artery obstruction, rheumatic disease, and the impossibility of undergoing the examinations. The patients in the chronic phase of the infection underwent complete clinical and cardiological examination and were classified as without demonstrable cardiac pathology or with cardiac involvement. The latter group was stratified according to modified Kuschnir classification (20).

**Table 1.** Demographic and clinical features of the study population.

| Characteristic | NI (Non *T. cruzi*-infected) | G0 (CCD without cardiac involvement) | G1 (CCD with cardiac involvement) |
| --- | --- | --- | --- |
| Age (Media and range) | 47 (30-60) | 57 (43-70) | 57 (44-74) |
| Gender (F/M) | 6/3 | 5/5 | 6/4 |
| Kuschnir Stages (0-1-2-3) | NA | 10-0-0-0 | 0-7-1-1 |
| PCR (+/-) | NA | 0/10 | 1/9 |
| Anti-*T. cruzi* IgG titre(Range)<sup>1</sup> | 10-32 | 1778-40439 | 6084-30318 |

<sup>1</sup>The titre of anti-*T.cruzi* antibodies was considered as IC<sub>50</sub> sera dilution factor. **CDD:** chronic Chagas disease.
**NA:** not applicable

An aliquot of whole blood (4 ml) from each participant was separated and centrifuged for 15 min at 800 g to obtain the plasma, which was stored at -20° until use. The rest of each sample was used to isolate PBMC. The titre of total anti-*T.cruzi* IgG antibodies in the plasma of patients was determined as previously described (21).

### Isolation and culture of PBMC

Peripheral blood mononuclear cells (PBMC) were isolated from whole blood by Ficoll-Hypaque density gradient centrifugation (GE Healthcare Bio-Sciences AB, Uppsala, Sweden) according to manufacturer-provided instructions, within 4 h after collection. Isolated PBMC were resuspended in fetal bovine serum (FBS, Natocor, Córdoba, Argentina) containing 10% dimethylsulphoxide and cryopreserved in liquid nitrogen until used. The number and viability of thawed PBMC were determined by Trypan blue exclusion staining in a Neubauer counting chamber. Cell suspensions were seeded in 48- well flat-bottom plates at a density of 2×10^6^ cells/well in 500 µl of RPMI-1640 medium supplemented with 100 U/ml penicillin, 100 µg/ml streptomycin, 2 mM L-glutamine and 10% heat-inactivated fetal bovine serum (FBS) at 37°C in a humidified 5% CO_2_ incubator. After 18 h of culture, cells were incubated for additional 5h with 50 ng/ml Phorbol-12- myristate-13-acetate (PMA) (InvivoGen, San Diego, CA, USA) + 1 µg/ml Ionomycin (MP Biomedicals, Santa Ana, CA, USA) or with culture medium only in the presence of 5¼g/ml Brefeldin A (Biolegend, San Diego, CA, USA) before being submitted to the flow cytometry staining process.

### Flow cytometry analysis of B and B10 cells

PBMC were transferred into a 96-well V-bottom plate and washed once with PBS by centrifugation at 700 g during 3 min at room temperature (RT). Supernatants were discarded and cells were resuspended in 25 µl of staining solution containing the antibodies detailed in **Table 2** diluted in 1X live/dead fixable viability dye (Zombie-Aqua, Biolegend) and incubated 30 min at RT in the dark. After surface markers staining, cells were washed with PBS, fixed with Fixation Buffer (Biolegend) for 20 min at RT in the dark and washed one more time with PBS by centrifugation at 700 g during 3 min at RT. For intracellular IL-10 detection, cells were permeabilized with Perm-Wash Buffer (Biolegend), stained with IL-10-PE mAb (**Table 2**) and fixed again with Fixation Buffer. “Fluorescence-minus-one” (FMO) controls were used to determine the cut point for the IL-10 staining, and isotype control staining was considered for cell surface markers gate setting. A minimum of 2×10^5^ events within the lymphocyte population were acquired in a FACSCanto II (BD Biosciences) flow cytometer using FACS Diva Software (BD Biosciences). Flow cytometry analysis was carried out with the program FlowJo (FlowJo LLC, Ashland, OR, USA). All antibodies were used at optimal concentrations determined by previous titration experiments. General gating strategy used to identify total lymphocytes population, B cells and CD4^+^/CD4^−^ T cells is detailed in **Sup. Fig.1** and subsequent gatings are described in representative dot plots in each Figure.

**Table 2.** Fluorescent-labeled antibodies and isotype controls used for flow cytometry experiments.

| **Antibody/Isotype control** | **Clone** | **Vendor** |
| --- | --- | --- |
| BV421- conjugated anti-CD3 | UCHT1 | Biolegend |
| PECy5- conjugated anti-CD4 | RPA-T4 | BD Biosciences |
| PECy5- conjugated anti-CD19 | HIB19 | BD Biosciences |
| APC- conjugated anti-CD27 | M-T271 | Biolegend |
| PECy7- conjugated anti-CD24 | ML5 | Biolegend |
| APCCy7- conjugated anti-CD38 | HB-7 | Biolegend |
| PE- conjugated anti-IL-10 | JES3-9D7 | Biolegend |
| BV421- mouse IgG1 $\kappa$ | MOPC-21 | Biolegend |
| PECy5- mouse IgG1 $\kappa$ | MOPC-21 | BD Biosciences |
| APC- mouse IgG1 $\kappa$ | MOPC-21 | BD Biosciences |
| PECy7- mouse IgG 2a $\kappa$ | MOPC-173 | Biolegend |
| APCCy7- mouse IgG1 $\kappa$ | MOPC-21 | Biolegend |

### Statistical analysis

To analyze the differences in frequency of cell populations between groups we applied a generalized linear model (GLM) (22) with quasi binomial distribution of errors and logit link function. Group and phenotype were considered fixed factors and the frequency of cells in each population was the dependent variable. Mean fluorescence intensity data were analyzed by a linear regression model (LM) fitted by maximum likelihood and Tukey HSD (honest significant difference) for post-hoc comparisons. The group was set as the fixed factor and mean fluorescence intensity was the dependent variable. MFI data were tested for normality and homoscedasticity using Shapiro-Wilk and Bartlett tests respectively. GLM and LM models were fitted in R3.5.1 (23). Results are represented with box and whiskers plots showing median value and interquartile ranges. *p-*values less than 0.05 were considered statistically significant.

## RESULTS

### Frequency of total and transitional B lymphocytes is altered in peripheral blood from patients with the cardiac form of chronic Chagas disease

In humans, four different subsets of peripheral blood B cells can be identified according to the expression levels of surface markers CD24 and CD38: CD19^+^CD24^high^CD38^high^ (immature transitional B cells), CD19^+^CD24^int^CD38^int^ (primarily mature naïve B cells), CD19^+^CD24^high^CD38^low^ (primarily memory B cells) and CD19^+^CD24^low^CD38^high^ (plasmablasts) (24, 25). Since the combined expression of these markers has not been addressed in the context of chronic *T. cruzi* infection, we decided to evaluate the frequency of the subsets mentioned above within total CD19^+^ B cells in PBMC from CCD patients without cardiac involvement (G0, n=10), CCD patients with cardiac involvement (G1, n=10) and non-infected donors (NI, n=9). First, we analyzed the frequency of total B lymphocytes and mean fluorescence intensity (MFI) for CD19 in CD19^+^cell population. The gating strategy used to identify total B cells (CD3^−^CD19^+^ cells) is illustrated in **Sup. Fig.1.** Results revealed higher percentage of CD19^+^B cells in patients with cardiac involvement compared to non-infected donors (**Fig. 1A**). No statistically significant differences were shown between G0 and NI or between CCD groups. Furthermore, similar MFI levels were found across the cohort, independently of the clinical status of the subjects (**Fig. 1B**).

**Figure 1:**
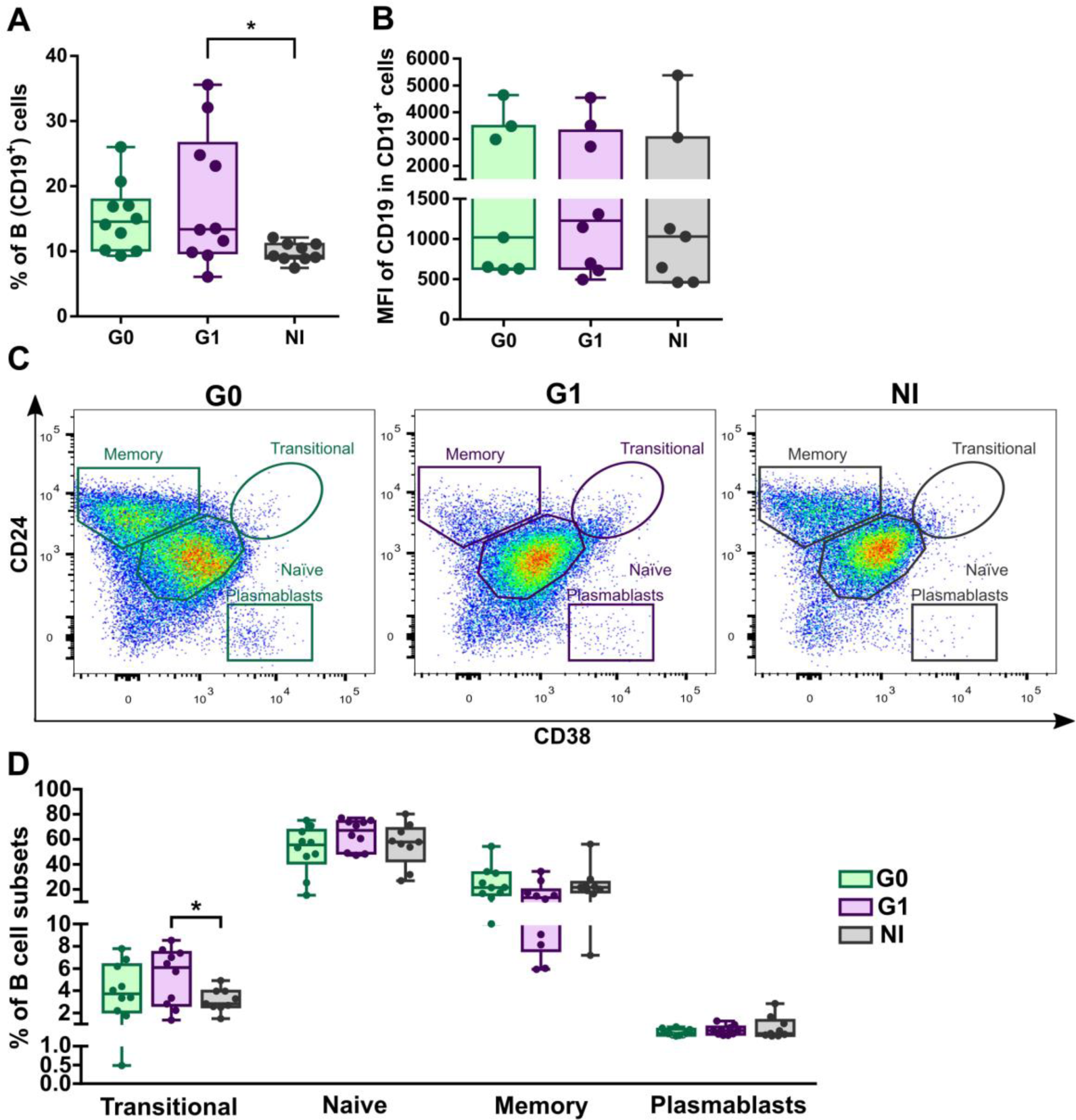
Frequency and phenotypic distribution of total B cells in peripheral blood from chronic Chagas patients and non-infected donors according to CD24 and CD38 expression. **(A)** Frequency of B cells (CD3^−^CD19^+^) and mean level of CD19 expression (as mean fluorescence intensity) in the CD3^−^CD19^+^ population from CCD patients (G0, G1, n=10) and non-infected donors (NI, n=9)**. (B)** Representative dot plots from CCD patients and one non-infected donor, showing the gating strategy used to identify B cell subsets. According to CD24 and CD38 expression levels B cells were sub gated into: CD19^+^ CD24^high^CD38^high^ (immature transitional B cells), CD19^+^ CD24^int^ CD38^int^ (mature naïve B cells), CD19^+^ CD24^high^CD38^−^ (Memory B cells), CD19^+^ CD24^low^ CD38^high^ (plasmablasts). **(C)** Frequency of different B cell subsets in total B cells from G0, G1 CCD patients and non-infected donors. Each dot represents data from one subject. Statistically significant differences among the groups are indicated (\**p*<0.05).

Next, using the gating strategy represented in **Fig. 1C**, we quantified the frequencies of different B cell subsets according to CD24 and CD38 expression in CCD patients and NI group. Results showed that the frequency of transitional B cells (CD24^high^CD38^high^) within total B cells were higher in patients with cardiac involvement from than those from non-infected donors. In addition, no significant differences were observed in the frequencies of the other B cell subpopulations among the groups (**Fig. 1D**).

Then, we determined the frequency of B cell subsets according to the expression of markers CD24 and CD27 or CD27 and CD38. The gating strategy to evaluate each subset of cells in a representative donor sample is illustrated in **Fig. 2A** and **Fig. 2B**. No differences among the groups were detected in any of the subpopulations evaluated (**Fig. 2C** and **Fig. 2D**).

**Figure 2:**
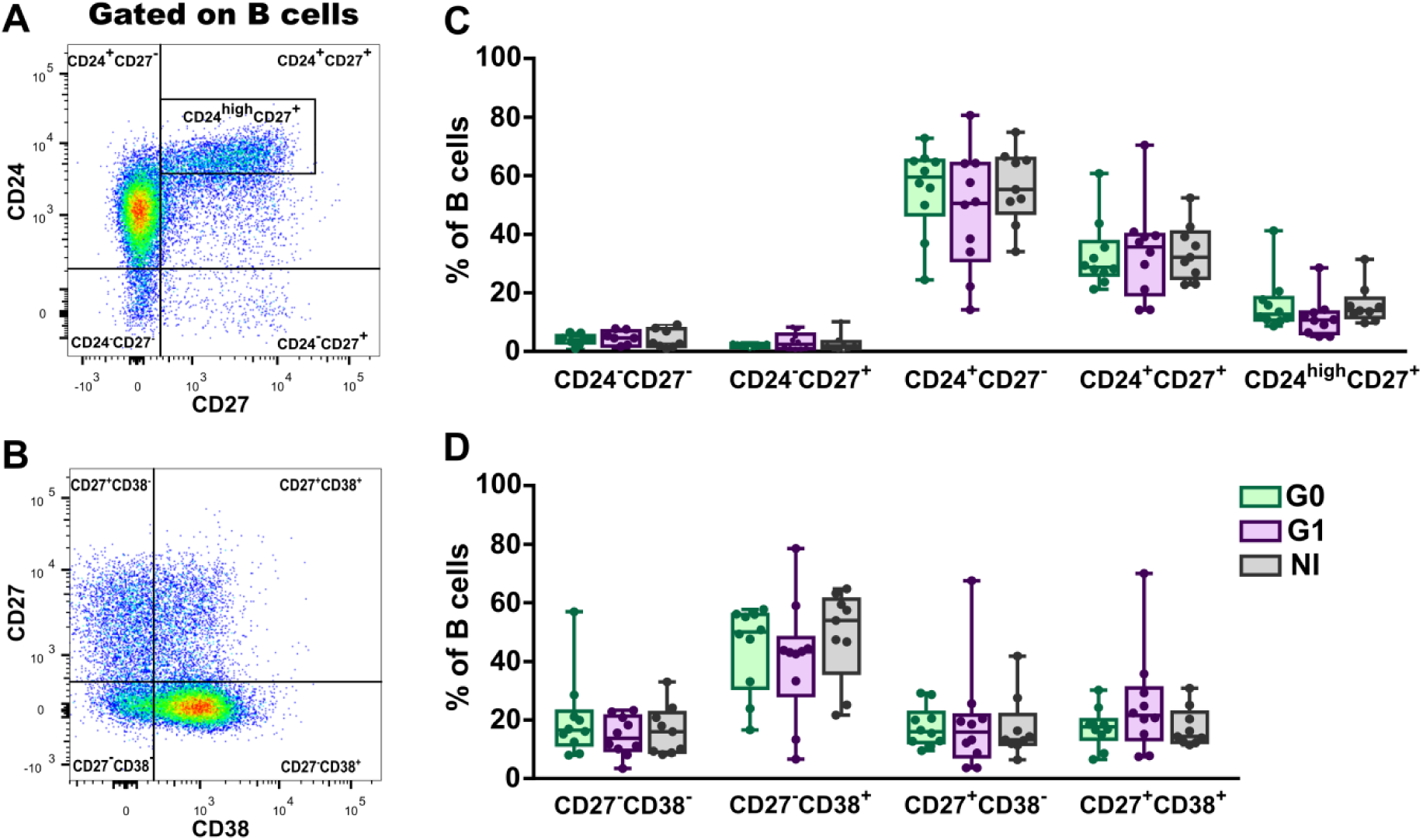
Frequency of B cell subsets according to CD24 and CD27 - CD27 and CD38 markers expression in peripheral blood from CCD patients and non-infected donors. **(A-B)** Representative dot plots from a non-infected donor showing the gating strategy used to determine subsets within total B cells according to the combined expression of CD24 and CD27 or CD27 and CD38. **(C-D)** Frequency of B cell subsets were evaluated according to CD24 and CD27 or CD27 and CD38 markers expression in CCD patients (G0, G1) and non-infected donors (NI). Each dot represents data from one subject.

### Phenotypic distribution but not frequency of the B10 cell compartment is altered in peripheral blood from patients with the cardiac form of CCD

Since CD24^high^CD38^high^ transitional B cell population has been associated with IL-10- mediated, regulatory B cell functions in autoimmune and infectious human diseases (11,26, 27), we next studied IL-10 producing B cells (B10 cells) in peripheral blood from patients with the different clinical forms of CCD. Thus, we performed an intracellular flow cytometry on PBMC, after *in vitro* incubation with PMA and ionomicyn in the presence of the protein transport inhibitor Brefeldin A for 5 h. We assessed the frequency of total B10 cells and B10 cell subsets according to CD24, CD38 and CD27 expression (11, 27) (**Fig. 3**). The gating strategy is shown in **Sup.Fig.1** and **Fig 3A**. Results showed that patients with the cardiac form (G1) have a tendency, although non-statistically significant to a decreased frequency of B10 cells compared to non-infected donors (**Fig 3B**). Furthermore, no differences in MFI of IL-10 in IL-10^+^ B cells were observed among the groups (**Fig. 3C**). The analysis of phenotypic distribution of B10 cells revealed that, for G0 and non-infected donors, most of B10 cells were contained within the naive (CD24^int^CD38^int^) and memory (CD24^high^CD38^low^) compartments, while a minor proportion was enclosed within the transitional subset (CD24^high^CD38^high^). In contrast, in patients with cardiac involvement (G1 group) B10 cells were distributed equally between the naïve and memory subsets, with a minor proportion belonging to the transitional compartment (**Fig. 3D**). In addition, the frequencies of B10 cells with transitional and naïve phenotype were higher in patients with cardiac involvement compared to those from non-infected donors (**Fig. 3D**). However, only the frequency of naïve B10 cells was statistically different among the groups (**Fig 3E**).

**Figure 3:**
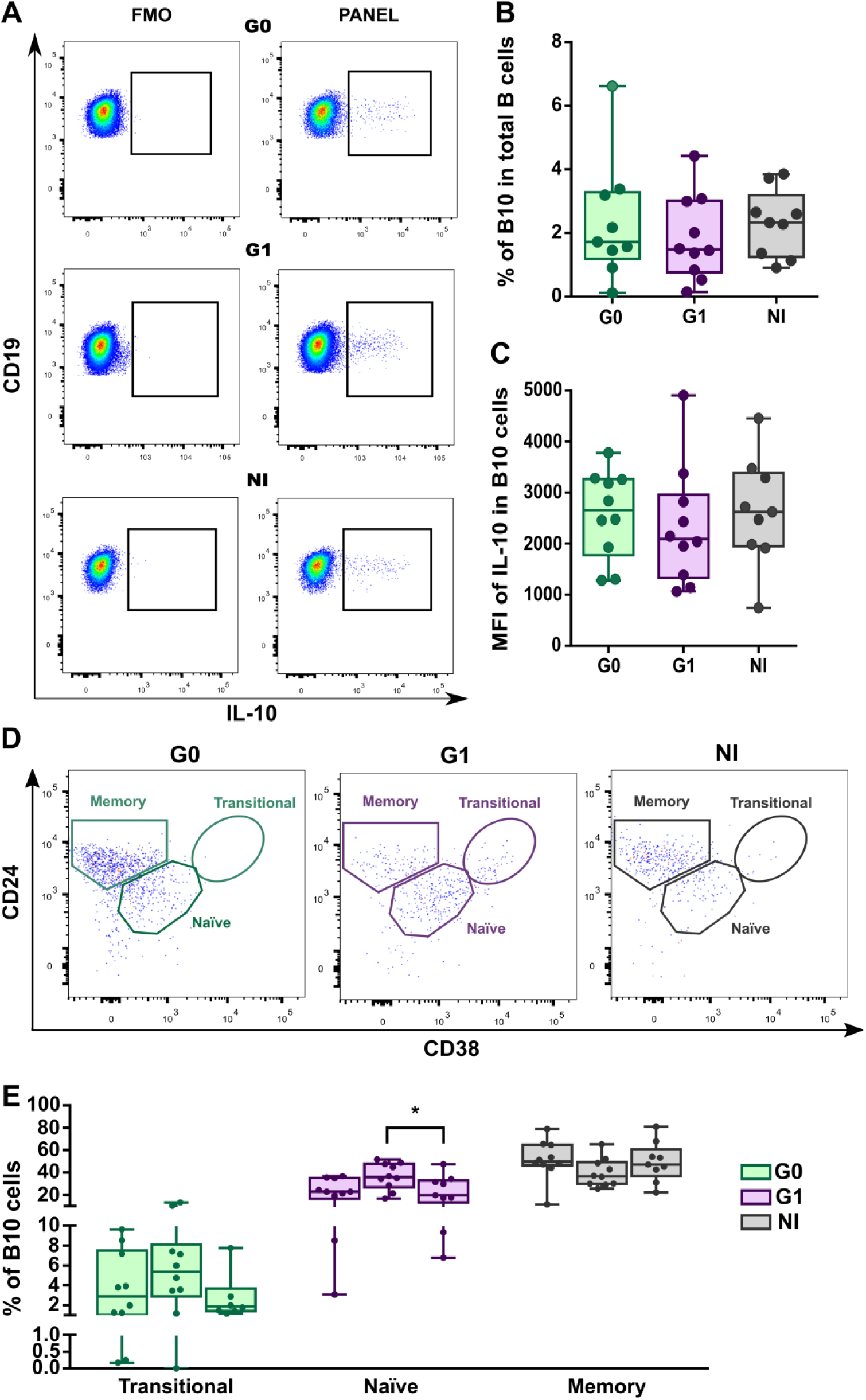
Frequency and phenotypic distribution of B10 cells according to CD24 and CD38 expression in peripheral blood from CCD patients and non-infected donors. **(A)** Representative dot plots showing the gating strategy used to determine IL-10^+^ B cells (B10 cells) in PMA + Iono stimulated PBMC. Fluorescence minus one controls were used to define negative populations for each sample. **(B)** Frequency of B10 cells in total B cells and mean fluorescence intensity for IL-10^+^ in B10 cells in CCD patients (G0, G1) and non-infected donors (NI). **(C)** Representative dot plots showing the gating strategy used to determine B10 cell subsets (transitional, naïve and memory) according to CD24 and CD38 expression in each group. IL-10 expressing B-cells (B10 cells) were gated according to subject- and condition-matched FMO control tubes, and were further sub-gated using CD24 and CD38 to identify the phenotypical distribution of these populations. **(D)** Frequency distribution of B10 cells according to CD24 and CD38 expression in CCD patients (G0, G1) and non-infected donors (NI). Each dot represents data from one. Statistically significant differences among subsets of cells and groups are indicated (\**p*<0.05).

It is well known that CD27 expression identifies human memory B cells (28). In order to better characterize the phenotypic distribution of B10 cells in CCD patients, we assessed the frequency of B10 cells according to CD27 expression alone and in combination with CD24 and CD38. According to CD27 expression, B10 cells were mainly found in the CD27^+^ (memory) population in the non-infected group, as was previously reported in samples from healthy blood donors (27) and the same pattern was observed in CCD patients (**Fig. 4B**). In addition, no differences were shown in the frequency of memory (CD27^+^) and naïve (CD27^−^) B10 cells subsets among the groups **(Fig. 4B**).

**Figure 4:**
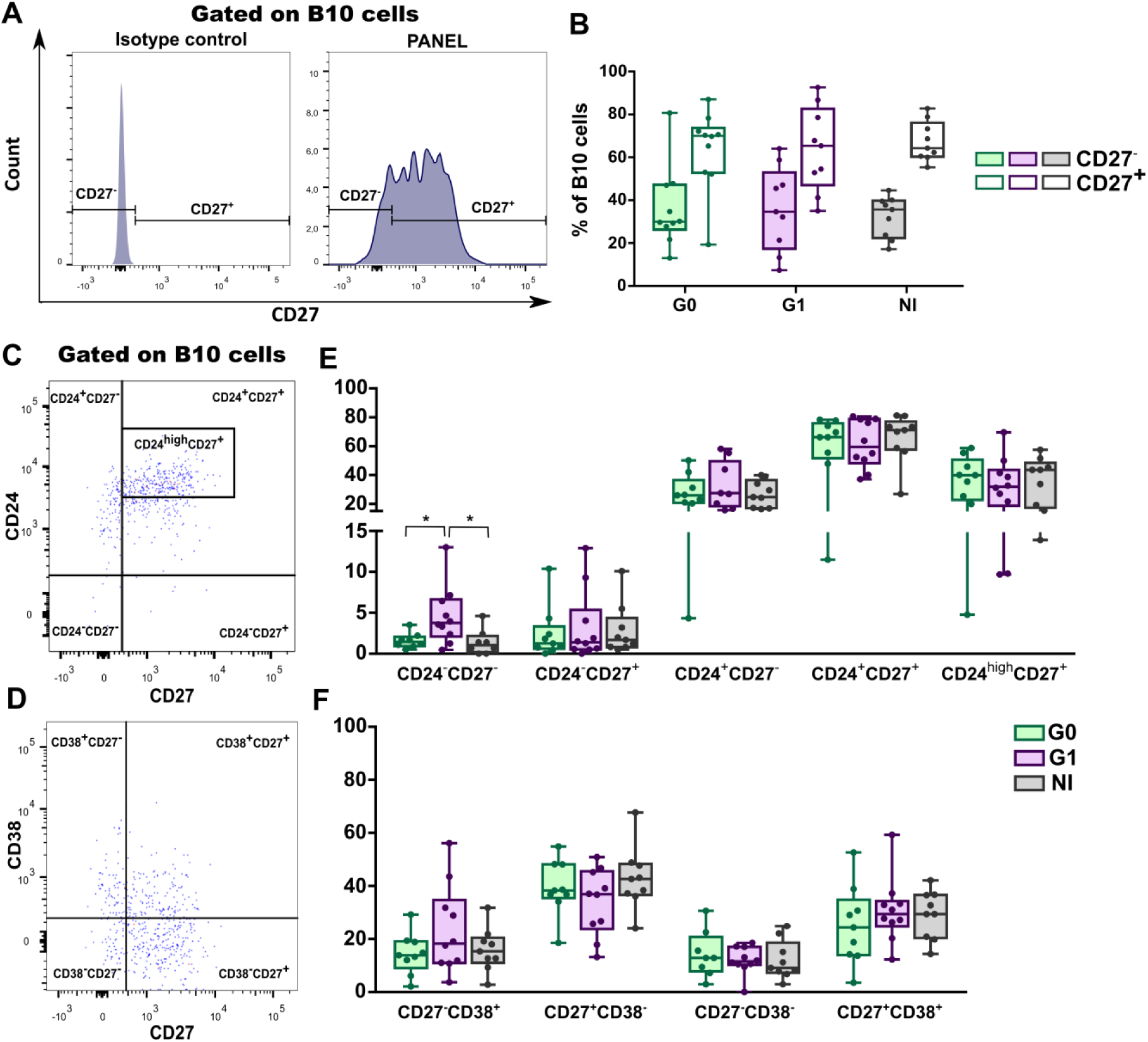
Phenotypic distribution of B10 cells according to CD27, CD27-CD24 and CD27-CD38 expression in CCD patients and non-infected donors. **(A)** Histograms showing the gating strategy used to determine CD27^−^ (naïve) and CD27^+^ (memory) populations within B10 cells in PMA + Iono stimulated PBMC. One representative non-infected donor is illustrated. **(B)** Frequency of CD27^−^ and CD27^+^ B10 cells in CCD patients (G0, G1) and non-infected donors (NI). **(C-D)** Representative dot plots from a non-infected donor showing gating strategy used to determine subsets within total B10 cells according to the combine expression of CD24 and CD27 or CD27 and CD38 in PMA + Iono stimulated PBMC. **(E-F)** Frequency of B10 cell subsets evaluated according to CD24-CD27 and CD27-CD38 markers expression in CCD patients (G0, G1) and non-infected donors (NI). Each dot represents data from one subject. Statistically significant differences among subsets of cells and groups are idicated (*\*p*<0.05).

Analysis of B10 cells subsets according to their CD24-CD27 expression showed an enrichment in CD24^−^CD27^−^ cells in patients with the cardiac clinical form compared to patients with no cardiac involvement (G0 group) and non-infected donors (**Fig. 4E**). Frequencies of B10 cell subsets according to CD24-CD38 expression did not shown statistically significant differences among the groups (**Fig. 4F**).

Given that IL-10 producing B cells are not confined to a single B cell subset, several studies have tried to determine which populations of B cell are enriched in IL-10^+^ cells. In this sense, most of the studies performed in human samples show that CD24^high^CD38^high^and CD24^high^CD27^+^compartments exhibit higher frequencies of IL-10 producing cells than their respective counterparts (11, 12, 27). Taking this into account, we aimed to evaluate the frequency of IL-10^+^cells within CD24^high^CD38^high^and CD24^high^CD27^+^ enriched subpopulations (**Fig. 5A** and **Fig. 5B** respectively). With this approach, we did not find differences between the groups (**Fig 5C** and **5D** respectively).

**Figure 5:**
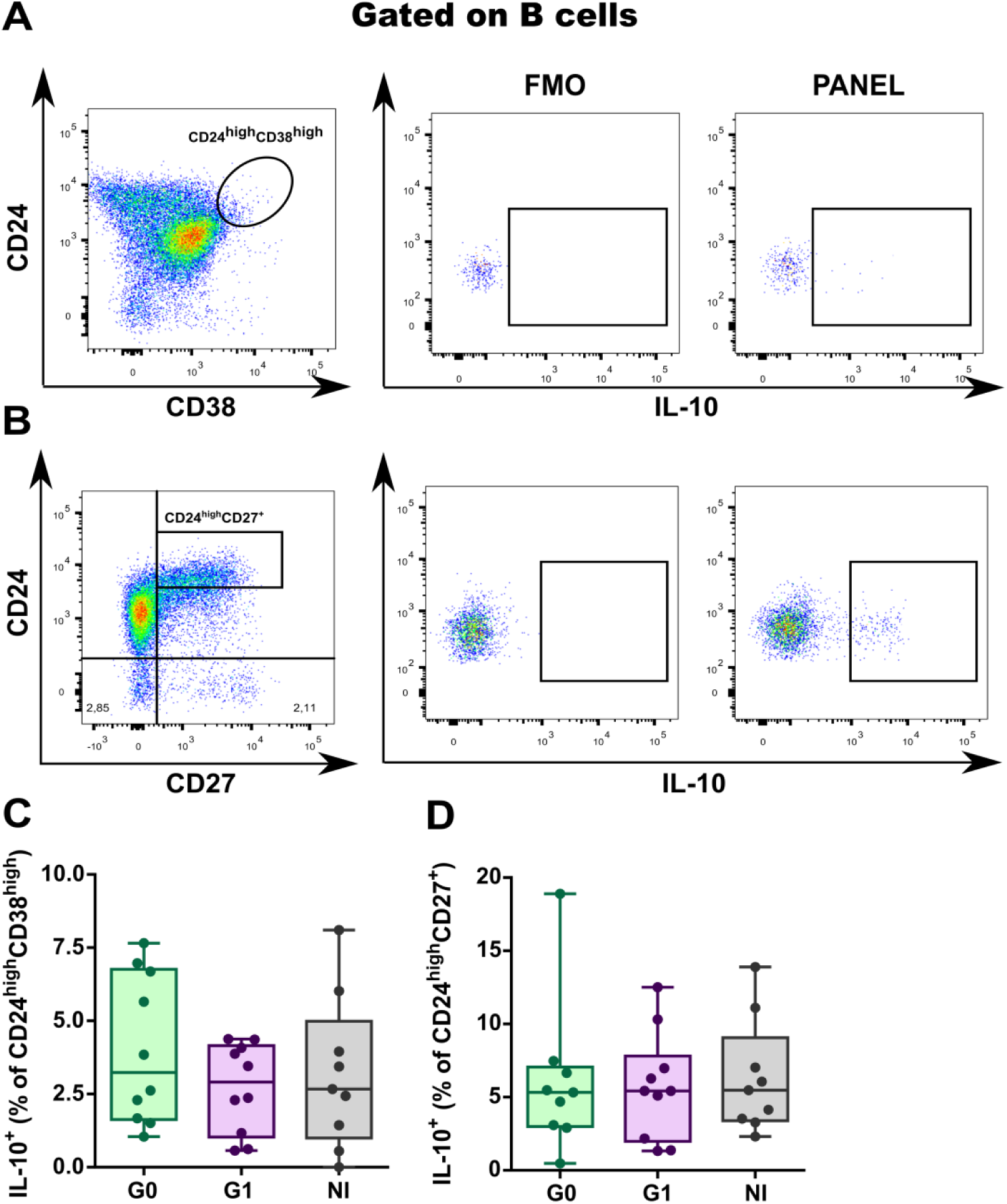
IL-10 producing cells within CD24^high^CD38^high^ transitional and CD24^high^CD27^+^ memory B cell populations in CCD patients and non-infected donors. **(A**, **B)** Representative dot plots showing the gating strategy used to determine frequency of IL-10^+^ B cells inCD24^high^CD38^high^ andCD24^high^CD27^+^B cells. **(C**, **D)** Frequencies of IL-10^+^cells within transitional (CD24^high^CD38^high^) and memory (CD24^high^CD27^+^) compartments in CCD patients (G0, G1) and non-infected donors (NI). Each dot represents data from one subject and boxes and whiskers show median value and interquartile range.

Production of IL-10 by PBMCs other than B cells showed that patients in G0 group tended to an increase in frequency of total IL-10^+^ lymphocytes (G0 vs. NI, *p*=0.056) and IL-10 was produced in comparable amounts by CD3^−^CD19^+^ (B cells) and CD3^+^ (T cells) in CCD patients and non-infected donors (**Fig. 6 A-C**). When analyzing inside the CD3^+^ population (**Fig. 6 D-G**), the frequency of IL-10^+^ CD4^−^T cells in CCD patients was increased compared to that in non-infected donors (**Fig. 6 F-G**).

**Figure 6:**
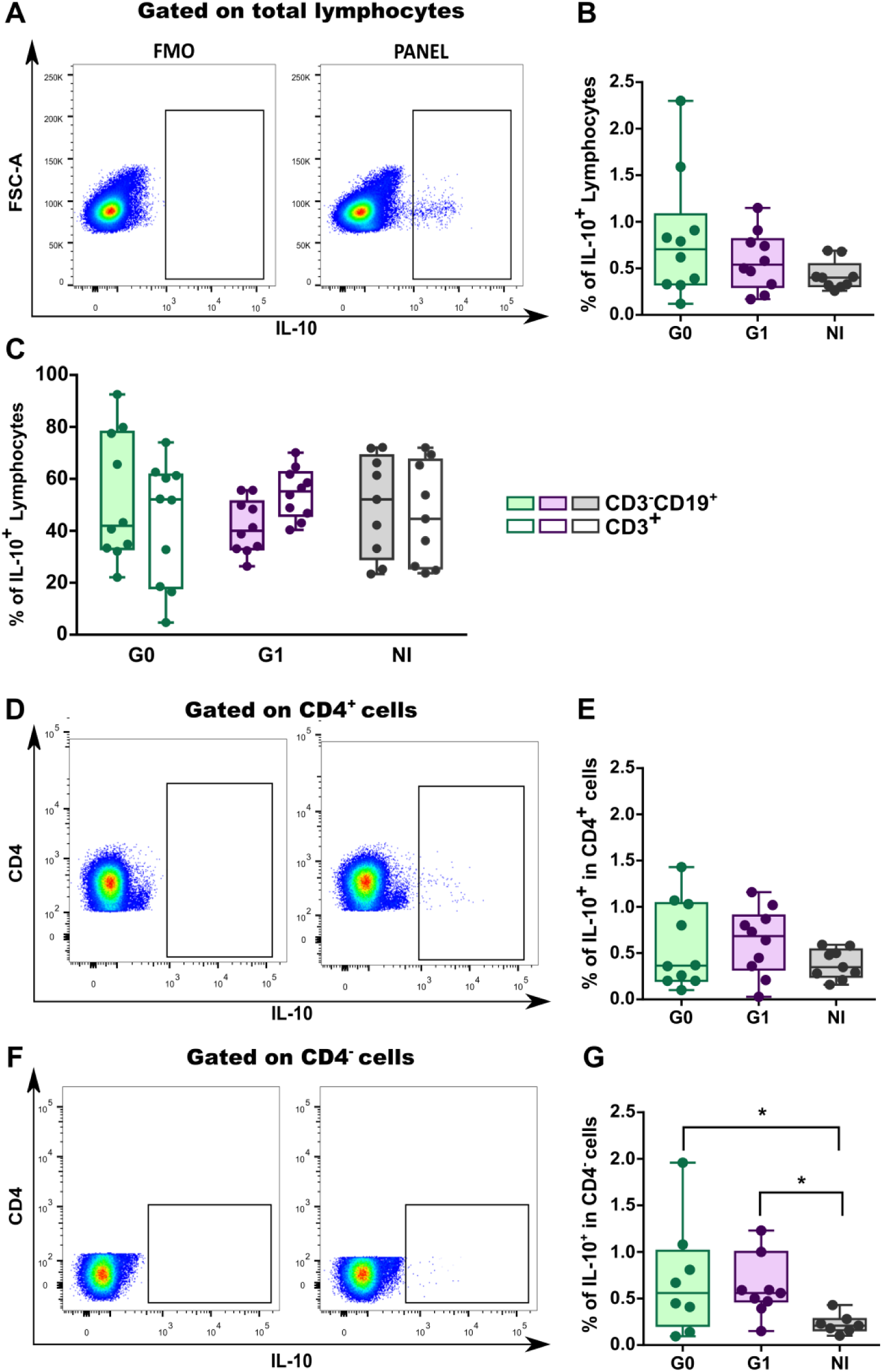
IL-10 production by total lymphocytes, B cells and CD4^+^/CD4^−^ T cells in CCD patients and non-infected donors. **(A-B)** Gating strategy and frequency of IL-10^+^ cells determined in total lymphocytes in CCD patients (G0, G1) and non-infected donors (NI). **(C)** Frequency of IL-10^+^ cells within B (CD3^−^CD19^+^) cells and T (CD3^+^) cells in PBMC from CCD patients (G0, G1) and non-infected donors (NI). Gating strategy employed is illustrated in Sup. Fig.1. **(D-G)** Frequency of IL-10^+^ cells within CD4^+^ and CD4^−^ T cell populations in CCD patients (G0, G1) and non-infected donors (NI). Each dot represents data from one subject. Statistically significant differences among groups are indicated (*\*p*<0.05).

## DISCUSSION

In this study, we characterized for the first time the B cell compartment in patients with chronic Chagas disease with and without cardiac involvement, in terms of their grouping in immature transitional (CD19^+^CD24^high^CD38^high^), primarily mature (CD19^+^CD24^int^CD38^int^), primarily memory (CD19^+^CD24^high^CD38^−^) B cells, and plasmablasts (CD19^+^CD24^−^CD38^high^) (24, 25).

We found that immature transitional peripheral blood B cells had higher frequencies in patients with cardiac involvement compared to non-infected subjects, and that this expansion could be conjoined with an increase in the percentage of total CD19^+^ B cells. In a previous work, Fernandez *et al* (2014) (29) observed an augmentation in the frequency of CD19^+^IgG^+^CD27^−^IgD^−^“double negative” memory and CD19^+^IgM^+^CD27^−^IgD^+^transitional/naïve B cells with a selective reduction of their classical memory CD27^+^ counterparts. In addition, and contrariwise to our observation, the frequency of CD19^+^ B cells in infected individuals was significantly lower compared to non-infected subjects. In this regard, and although it would be interesting to correlate our findings with those described therein, it should be noted that B cell phenotypes were designated on the basis of different surface markers. In addition, the infected subjects in the cohort of the mentioned study were grouped together for the analysis, independently of their clinical status. However, both studies clearly depict an alteration in the peripheral B cell compartments of infected individuals during the chronic phase of Chagas disease with a predominance of intermediate forms between the immature and mature phenotypes. This phenomenon could be due to genetically intrinsic B cell abnormalities as well as extrinsic cellular or molecular factors that regulate B cell lymphopoiesis (30). Accordingly, there are evidences that BAFF the B-cell activating factor of the TNF family which is essential for transitional B cells to outright their development (31-34) and myeloid cell derived factors are abundantly secreted by the parasite early after infection, leading not only to interferences in mature B cell development but also to an autoimmune response through polyclonal activation (35-38). Besides, another important checkpoint necessary for the development of human B cell precursors is the IL-7/IL-7R axis (39) which was found to be altered in chronic Chagas disease patients with severe cardiomyopathy (40).

Production of IL-10 is the ultimate feature of several described B regulatory phenotypes (41), and is the broadest phenotypic marker of human Bregs described so far (42). Several subsets of IL-10 producing B cells have been described in humans, their definition depending on the disease under study, the immunological context and the stimulation conditions. These include CD1d^+^CD5^+^ (43, 44), Tim-1^+^(15, 16), immature transitional CD24^hi^CD38^hi^ (11, 41) and memory CD24^high^CD27^+^ B cells (12).

Focusing on chronic infections, a potent immunosuppressive role has been reported for B10 cells, acting mainly upon antigen-specific CD8^+^ T cells (45). In HIV-infected children and adolescents, an increased frequency of circulating Breg cells induced by influenza immunization has been linked to a poor response to vaccination (46). In leishmaniasis, IL- 10 secreted by CD5^+^CD1d^+^ B cells polarized the Th cell response toward the Th2 phenotype, necessary for susceptibility to infection with the parasite in BALB/c mice (47). Even more, incubation of human B cells with *Leishmania infantum* amastigotes induced IL- 10 production by CD24^+^CD27^−^ B cells that modulated partially INF-γ and TNF-α secretion by T cells (48). Although B10 cells are involved in the maintenance of homeostasis in the immune system (5, 49), these findings highlight the importance of the immunosuppressive mechanisms triggered by microorganisms which could favor their persistence and contribute to poor vaccine response (50, 51). Nonetheless, and given the implications of uncontrolled inflammation in the context of Chagas disease, B10 cells are likely to contribute to the delicate balance between parasite clearance and inflammatory pathogenesis.

As mentioned above, the only work attempting to analyze the role of B10 cells in chronic Chagas disease has been carried out by Fares et al (2013)(19). Therein, *T. cruzi* infected patients, independently of the clinical form of the disease, presented a slightly higher frequency of IL-10-producing CD19^+^CD1d^+^CD5^+^ cells compared to non-infected subjects. Here, we found that the frequency of B10 cells was similar among the groups, but the phenotypic distribution based on CD24 and CD38 surface markers was altered in patients with cardiac involvement compared to patients without cardiac involvement and non-infected subjects. While no differences were observed in the frequency and phenotypic distribution of B10 cells between non-infected and patients without cardiac involvement, those with cardiac manifestations had significantly larger naïve and immature transitional subpopulations within B10 cells. This augmentation seems to be associated with a decrease tendency in the frequency of CD24^high^CD38^low^B10 cells. Interestingly, although, the other phenotype associated with Breg cells CD24^high^CD27^+^ IL-10 producing cells, seemed not to be altered, the percentage of CD24^−^CD27^−^ cells was also significantly increased in patients with cardiac manifestations. The functional features of this subpopulation remain elusive and the relevance of this augmentation will be a matter of further investigation in our laboratory.

Because IL-10 producing cells within CD24^high^CD38^high^B cell and CD24^high^CD27^+^ B cells have been broadly linked to immune regulation and control of chronic inflammatory diseases, autoimmunity and graft *vs*. host disease (11, 12, 17, 27) we decided to assess IL-10 production in these B cell subsets. Our data revealed no differences among the individuals independently of their clinical status, suggesting that B cells from patients with cardiac involvement have a regulatory phenotype mainly due to an expansion in CD24^high^CD38^high^ immature transitional compartment.

Chronic Chagas cardiomyopathy and dilated cardiomyopathy (DCM) have a similar structural disarrangement that leads to ventricular dilatation, but the former is characterized by severe myocarditis and a dense fibrosis that surround each myocardial fiber or group of myocardial fibers (52). Conversely to our observation, YujieGuo et al (2015)(53) showed that the frequency of CD19^+^IL-10^+^ B cells in peripheral blood from DCM patients was higher than those from healthy individuals under the same stimulation conditions followed in our work. However, when stimulated with CD40L and CpG, the percentage of CD19^+^IL-10^+^ B cells diminished in DCM patients compared to a healthy control group (54). In addition, the frequency of CD24^high^CD38^high^ transitional B cells and CD24^high^CD27^+^ B cells in DCM patients were comparable with that of healthy individuals, while IL-10 production of CD24^hi^CD27^+^ B cells from DCM patients was decreased. Although the experimental conditions differ between our study and that by Jiao et al 2018 (54) and this could account for the discrepancies seen in the populations mentioned above, we cannot rule out that these alterations may be responsible of the histopathological findings ascribed to each disease.

Human CD24^high^CD38^high^ immature transitional B cells from healthy individuals seem to inhibit T cell polarization towards Th1 and Th17 profiles while prompting the conversion of CD4^+^CD25^−^ T cells into Treg cells, mainly through IL-10 release (11). However, their function shifts in immune disorders, like in rheumatoid arthritis where transitional B cells fail to convert CD4^+^CD25^−^ T cells into functionally suppressive Treg cells or to restrain Th17 development, albeit maintaining the capacity to inhibit Th1 cell differentiation (55). This differential activity may be related with the presence of two subsets of CD19^+^CD24^hi^CD38^hi^B cells, the functionally ‘‘conventional’’ immature B cells programmed to become mature B cells and the B cells that are functionally regulatory (11). Notably, the capacity of impairing conversion of T cells towards a Treg cell profile is not an exclusive feature of CD19^+^CD24^hi^CD38^hi^ B cells. In DCM, CD24^hi^CD27^+^B cells have a decreased ability to suppress the TNF-α production by CD4^+^CD25^−^T cells and to maintain Treg cells differentiation, leading to worsen the cardiac condition (54).

In summary, our work shows that immature transitional CD24^high^CD38^high^ B cells are greatly expanded in patients with the cardiac form of chronic Chagas disease, and that these cells do not lose their capacity to secrete IL-10 compared to non-infected donors. This expansion could be the result of defective B cell development during acute *T. cruzi* infection or may be the consequence of the pro-inflammatory state. Remarkably the distribution of naïve, transitional and memory B cells inside the B10 cells follow the same pattern in chronic patients without cardiac involvement and non-infected individuals. Finally, our data are insufficient to define whether these immature transitional CD24^high^CD38^high^ B cells have deficiencies in their regulatory function and correlate with the clinical outcome of patients with chronic Chagas disease. We expect that our ongoing *in vitro* studies aiming at the elucidation of the Breg cells’ impact on CD4^+^ and CD8^+^ T cells may help to address this matter.

## ACKNOWLEDGEMENTS

This work was supported by the Consejo Nacional de Investigaciones Científicas y Tecnológicas (CONICET; Grant number 112-200801-02915) and by the Agencia Nacional de Promoción Científica y Tecnológica (ANPCyT; Grant number 2014-1026) from Karina. A Gómez and by Fundación AJ. ROEMMERS from Gonzalo R. Acevedo. We thank all the patients and non-infected individuals who participated in this study. We are also very grateful to Violeta Chiauzzi from Instituto de Biología y Medicina Experimental (IBYME-CONICET) for technical assistance in blood sample collection.

