## Supplementary Figure 1 for "Altered frequency of CD24^high^CD38^high^ transitional B cells in patients with cardiac involvement of chronic Chagas disease"

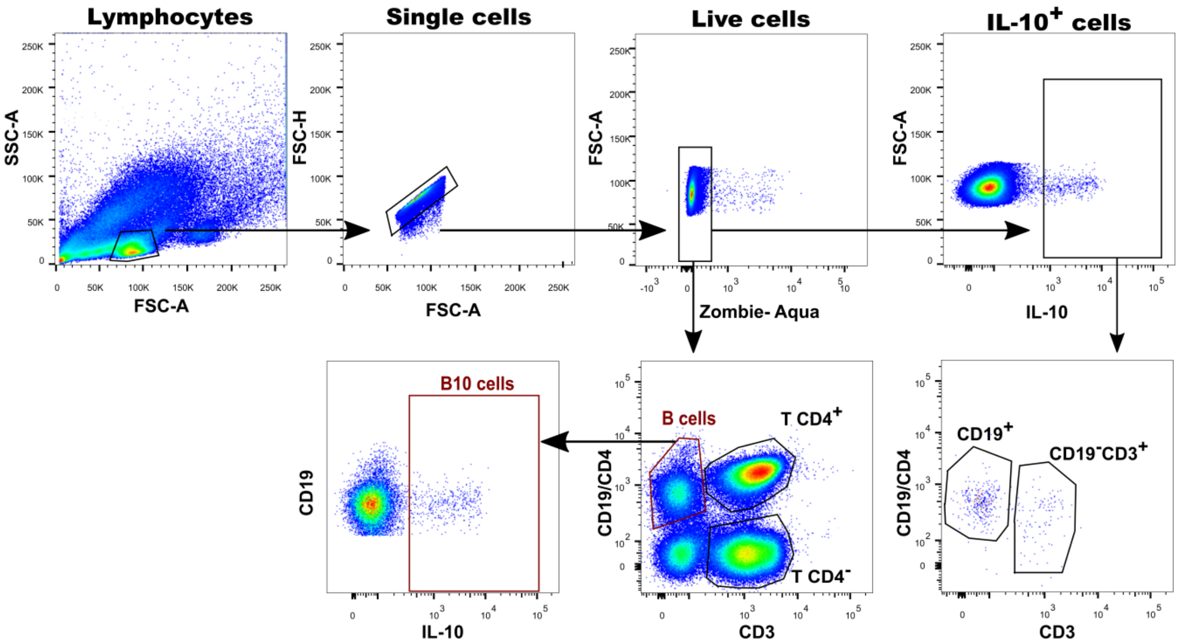


**Supplementary Fig. 1: General gating strategy used to define the lymphocyte populations to be studied in CCD patients and non-infected donors.** Representative plots obtained from the analysis of a sample by flow cytometry, according to the expression of CD3, CD4, CD19 and IL-10 markers. Cells were gated according to forward scatter/side scatter area (FSC-A/SSC-A) criteria to identify the lymphocytes population. Non-single cell events and cells with compromised viability were excluded by gating on the FSC-A/FSC-H and Zombie Aqua channels, respectively. B-cells were gated using CD19 and CD3 markers ($CD3^{-}CD19^{+}$ population). Within viable lymphocytes, total IL-$10^{+}$ cells were determined and then the frequency distribution of total IL-$10^{+}$ cells within $CD3^{-}CD19^{+}$ (B cells) and $CD3^{+}$ (T cells) populations was evaluated. Isotype controls were used to determine cut points for each marker. IL-$10^{+}$ cells were gated according to subject- and condition-matched FMO control tubes. B and B10 cells were further sub-gated according to CD24, CD38 and CD27 markers to identify the phenotypical distribution *ex vivo* of these populations. Further gating analysis is shown in the corresponding figures. Data analysis was performed using the program FlowJo.
